# Iron dysregulation in mice engineered with a mutation associated with stuttering

**DOI:** 10.1101/2025.07.30.667752

**Authors:** Marissa Millwater, Camryn Bragg, Devin Bishop, Afuh Adeck, Rahul Chowdary Karutury, Maximillian Weinhold, Praveen P.N. Rao, Ruli Zhang, Shahriar SheikhBahaei

## Abstract

Stuttering is a neurodevelopmental disorder characterized by involuntary disruptions in the normal fluency and timing of speech. Recently, stuttering has been related to specific point mutations in *GNPTAB*, a gene involved in lysosomal enzyme-targeting pathways, though it remains unclear how such a mutation might cause the stuttering phenotype. Herein, we studied mice engineered with the mutation in the *Gnptab* gene found in humans who stutter and found increased iron deposition in the basal ganglia of these mice. Further, we found these iron deposits localized predominantly with regional astrocytes when Perls’ stain was combined with an astrocyte-specific marker. Reducing iron deposition in the brain with iron chelation therapy improved vocalization symptoms in *Gnptab*-mutant mice. Our data suggest a relationship between the *Gnptab* mutation, iron homeostasis in astrocytes, and the stuttering phenotype, for which the underlying mechanisms remain to be elucidated.

## 1 INTRODUCTION

It is estimated that more than 80 million adults worldwide are affected by stuttering, a neurodevelopmental disorder characterized by involuntary disruptions in speech fluency. Developmental stuttering, also known as childhood-onset fluency disorder, emerges in early childhood, and while its exact cause(s) is not fully understood, it is believed to involve a combination of genetic, neurological, and environmental factors (SheikhBahaei et al., 2023). Current data suggests that stuttering is associated with subtle differences in brain structure and function, particularly in areas related to speech production, but it does not involve overt brain damage nor widespread disruptions to other brain functions (Chang et al., 2008; Watkins et al., 2008. Associations have been made between stuttering and dopaminergic hyperactivity, cortical connectivity impairment, and white matter anomalies in the brain (Maguire et al., 2021; Chang et al., 2019; Cieslak et al., 2015). More recently, iron accumulation has been reported in adults who stutter (Cler et al., 2021; Liman et al., 2021).

Iron is a necessary transition metal implicated in mitochondrial function, myelin production, and the synthesis of neurotransmitters (Ward et al., 2014; Cheli et al., 2020). However, whether iron is to the benefit or detriment of cellular processes and larger organ systems is dependent on its proper regulation and homeostasis (Morris et al., 1992; McCarthy et al., 2015). Recent transcranial ultrasounds of the midbrain detected moderate mesencephalic hyperechogenicity, which is likely associated with iron deposits, in ten adults who stuttered since childhood versus ten adult controls (Cler et al., 2021; Liman et al., 2021).

It was previously determined that mutations in the N-acetylglucosamine-1-phosphate transferase gene (*GNPTAB*) and three other related genes (*i*.*e*., *GNPTG, NAGPA*, and *AP4E*) are linked to developmental stuttering (Kang et al., 2010; Drayna et al., 2017). The human *GNPTAB* stuttering mutations (Glu1200Lys) were since engineered into mice that display vocal deficit (Barnes et al., 2016; Han et al., 2019). However, it was unknown if these mice show increased iron levels in the brain.

In the current study, we aimed to confirm, assess, and alleviate iron accumulation in the knock-in transgenic mouse engineered with the mutation in the *Gnptab* gene. Our data suggest an increase in brain iron levels in *Gnptab*-mutant mice compared to control littermate adult mice in striatum, a basal ganglia region related to control of motor behaviors. In addition, we identified changes in morphology of astrocytes, the ubiquitous star-shaped glia cells in the brain, in the striatum. Treatment of *Gnptab*-mutant mice with an iron chelator drug, deferiprone, improved the vocal deficits in the mutant mice. Taken together, our data suggest a potential mechanistic link between a mutation in a lysosomal trafficking gene and increased iron concentration in brain regions that are linked to stuttering disorders.

## 2 METHODS

### 2.1 Animals

All procedures in this study were performed in accordance with Guide for the Care and Use of Laboratory Animals and with Institutional Animal Care and Use Committee approvals National Academies Press (US), 2011). Male mice were group-housed in cages with a 12 h light/dark cycle unless otherwise stated. Mice were maintained *ad libitum* on a standard diet and water.

### 2.2 Vocalization studies

To assess social vocalization patterns, *Gnptab*-mutant (n = 11) and control (n = 10) male mice were placed in a clean testing environment and allowed to acclimate for 20 minutes. A female mouse was introduced for 5 minutes, allowing for the interaction of the male and female mice. The female was removed, and the male was allowed to re-acclimate to the existing environment. During this reacclimating period, an absorbent pad was placed under the bedding of the female cage to collect urine and other debris. The absorbent pad was subsequently placed in the testing environment to induce vocalization, which was recorded for 3 minutes by an Avisoft ultrasonic microphone and via Avisoft-RECORDER (RRID: SCR_014436; Avisoft Bioacoustics). For vocalization classification, WAV files were imported to Sonotrack Call Classification. The classifications were exported to Prism for statistical analysis.

### 2.3 Iron chelation therapy

*Gnptab*-mutant and control littermates from the pre-treatment studies described above were exposed to an iron chelation therapy for 1 month via the water supply. The iron chelator drug deferiprone, 3-hydroxy-1,2-dimethylpyridin-4-one (DFP), was dissolved in animal water (0.1mg/L) and replaced as needed. No health changes were observed throughout the course of treatment, as reported elsewhere (Devos et al., 2014; Martin-Bastida et al., 2017; Devos et al., 2022). After one month of iron chelator therapy, vocal behaviors of all treated mice were reassessed. Three mice per group were chosen for histological analysis (see below). Other animals were placed on normal water for 30 days after which the vocal behaviors were re-assessed.

### 2.4 Tissue processing and histological processing

At the completion of the experiment, mice were transcardially perfused with 50 mL of 0.9% saline solution followed by 4% paraformaldehyde (PFA) fixative, as described before (Hosford et al., 2020). The brain was extracted and post-fixed for 2-3 days in the same PFA solution. The extracted brains were sectioned at 50•m. Floating slices were stored in an anti-freeze solution at –20 ºC until staining. Sections were selected based on the corresponding section in the mouse brain atlas (Paxinos and Franklin et al., 2019).

Iron was detected by diaminobenzidine-enhanced (DAB-enhanced) Perls’ stain (Turnbull method), from here on referred to as “Perls’ stain,”, which stains iron in the ferric state, including ferritin and hemosiderin (Meguro et al., 2007; Lee et al., 2019). Optimal Perls’ staining was observed when slices were chemically stained within 3 weeks of perfusion to avoid loss of iron to the storage solution over time. Further, the aggregation of iron deposits near regional neurons and glial cells (i.e. astrocytes, oligodendrocytes, microglia) was investigated by combined immunohistochemistry and Perls’ staining. Antibodies for these immunohistochemical analyses are detailed in Table 1. Perls’-stained regions of interest were imaged via bright field microscopy (Zeiss Axioscan) with a 20x objective. Tiled or single images were imported to ImageJ-FIJI, where the percentage of iron-stained area was quantified.

**Table 1.**
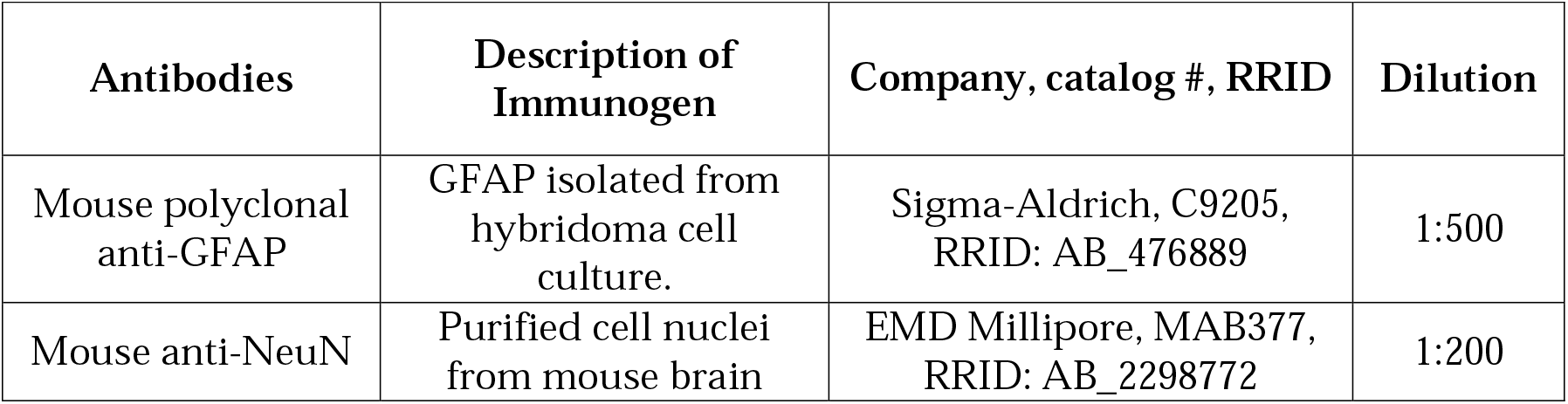
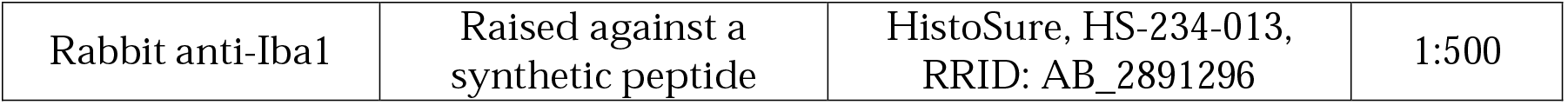
Primary antibody characterization.

| Antibodies | Description of Immunogen | Company, catalog #, RRID | Dilution |
| --- | --- | --- | --- |
| Mouse polyclonal anti-GFAP | GFAP isolated from hybridoma cell culture. | Sigma-Aldrich, C9205, RRID: AB_476889 | 1:500 |
| Mouse anti-NeuN | Purified cell nuclei from mouse brain | EMD Millipore, MAB377, RRID: AB_2298772 | 1:200 |
| Rabbit anti-Iba1 | Raised against a synthetic peptide | HistoSure, HS-234-013, RRID: AB_2891296 | 1:500 |

**Table 2.** Secondary antibody characterization (Perls’ staining)

| Antibodies | Source, host species, catalog no., RRID |
| --- | --- |
| ImmPRESS – AP Horse Anti-Mouse IgG Polymer Kit, Alkaline Phosphatase | Vector Laboratories, MP-5402, RRID: AB_2336535 |
| ImmPRESS – AP Horse Anti-Rabbit IgG Polymer Kit, Alkaline Phosphatase | Vector Laboratories, MP-5401, RRID: AB_2336536 |

### 2.5 Reconstruction and morphometric analysis of astrocytes

For morphometric analysis, astrocytes were immunostained with anti-GFAP antibody as described before (SheikhBahaei et al., 2018; Adeck et al., 2024). Confocal image stacks of the GFAP-positive astrocytes from the striatum were taken via an inverted laser scanning microscope (Zeiss LSM 510) using a high magnification, oil immersion objective (40×/1.2 NA) with 1024•×•1024 pixels resolution (SheikhBahaei et al., 2018). Confocal z-stack images were imported into Imaris 9.7.0 (Oxford Instruments, RRID: SCR_007370), where reconstructions of individual astrocytes were performed with Imaris software tracing tools as described before (Turk et al., 2022). Briefly, images were optimally obtained to cover regions of interest and minimize potential overlap from other regions. GFAP-positive astrocytes with fully intact processes from the striatum were selected for reconstruction (five to eight images across the rostral-caudal axis; two to four astrocytes per image). Astrocytic processes were traced by one investigator and verified by a second investigator.

#### Morphometric analysis of astrocytes

Imaris was used to process and retrieve morphometric data from fully traced and reconstructed astrocytes as described before (Turk and SheikhBahaei et al., 2022; Adeck et al., 2024). The 3D filament data tool was utilized to carry out morphometric analysis of astrocytes in regions of interest. Extracted features included Sholl analysis (Sholl et al., 1953) to determine the unique cellular process found in each region. Sholl analysis was performed to account for the variation in complexity of astrocytes over the radial distance from the soma by quantifying process branches, branch points, number of terminal points, as well as process length in astrocytes. This analysis utilized shell volumes between concentric spheres, each 1•m apart, radiating out from the center of the soma. Because of the complexity of astrocyte morphology, the 3D convex hull analysis was used to estimate the volume of astrocytes as described before (Sheikhbahaei et al., 2018; Turk and SheikhBahaei et al., 2022; Adeck et al., 2024). In convex hull analysis, the volume and surface area of astrocytes were estimated by enveloping the cell surface area and volume, creating a polygon that joins terminal points of the processes. We then used the complexity index (CI) to normalize the comparison of the overall cellular complexity in distinct regions as previously described (SheikhBahaei et al., 2018; Turk and SheikhBahaei et al., 2022; Adeck et al., 2024). The CI has also been used to evaluate the complexity of neurons (Diamantaki et al., 2016; Guillamon-Vivancos et al., 2019; Reagan et al., 2021) and glia cells (Sheikhbahaei et al., 2018a; Turk and SheikhBahaei, 2022).

### 2.6 Statistical analysis

The analyzed morphometric data and USVs were exported from Imaris to Prism 10 software (Graphpad Software Inc., RRID: SCR_002798) for statistical analysis. The data were reported as averages ± standard error of the mean (sem) and graphed as box and whisker plots. An unpaired *t-*test was used for statistical analysis.

## 3 RESULTS

### 3.1 Vocalization phenotypes of adult Gnptab-mutant mice

Previous data reported that neonate mice carrying a homozygous Glu1179Lys mutation in the *Gnptab* gene (*Gnptab*-mutant) emit fewer isolation calls and increased pause length between vocalizations (Barnes et al., 2016). However, the vocal behaviors of adult *Gnptab*-mutant animals have not been fully evaluated. Therefore, we first analyzed vocal behaviors in adult male mice. As opposed to pup calls, we did not find any differences in frequency of vocalizations in adult animals (201 ± 19 min^-1^ vs 187 ± 24 min^-1^ in control; *P* = 0.7, unpaired *t*-test; Figure 1). However, the geometry (spectral characteristics) of vocalizations was affected by the mutation, as the normalized ratio of compound vocalization was higher in the *Gnptab*-mutant compared to those in control littermates (0.15 ± 0.01 vs 0.10 ± 0.01 in control; *P* <0.001, unpaired *t*-test; Figure 1).

**Figure 1.**
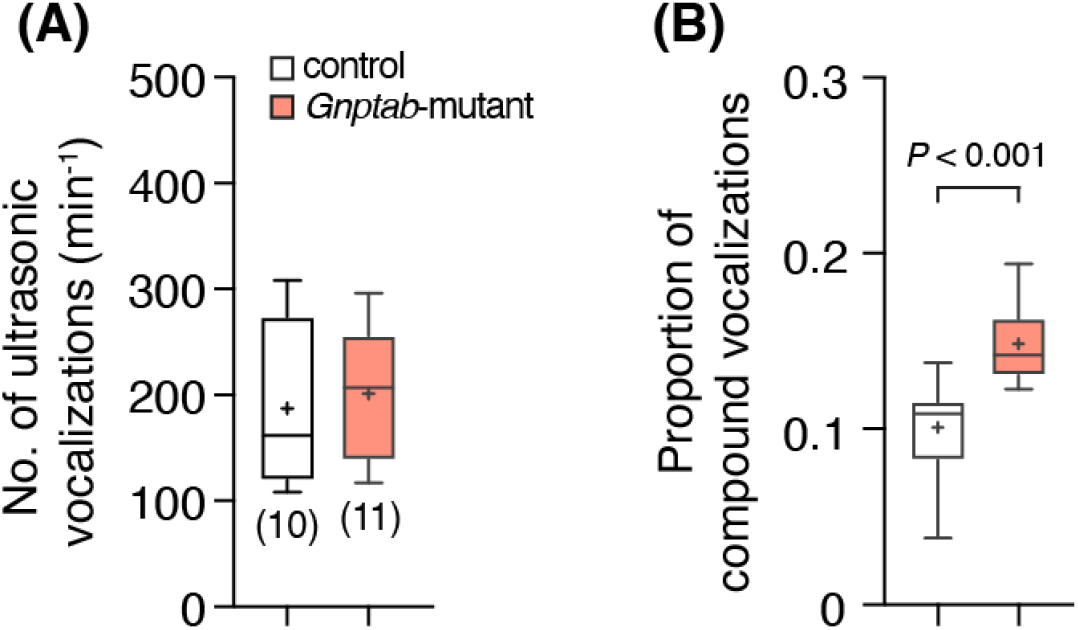
*Gnptab*-mutant male mice showed a higher frequency of compound vocalization. Frequency of vocalizations (calls per minute) in adult *Gnptab*-mutant mice (n = 11) and control littermates (n = 10) shows no significant difference (*P* = 0.7, unpaired *t*-test). **(B)** Summary data indicating changes in spectral characteristics of ultrasonic vocalizations, represented by an increase in the ratio of compound vocalizations in *Gnptab-*mutant mice compared to controls (*P* – *t*-test unpaired *t*-test).

### 3.2 Increased iron concentration in the striatum of Gnptab-mutant mice compared to control littermates

Since higher concentration of iron is reported in the brains of adults who stutter (Cler et al., 2021; Liman et al., 2021), we next assessed whether iron also accumulates in *Gnptab*-mutant animals. Our data directly showed higher iron concentration, detected by Perls’ staining, in the brain of *Gnptab*-mutant mice (Figure 2). The iron deposition was more prominent in the striatum (the largest structure of the subcortical basal ganglia). Our data suggest an increase in iron levels in the striatum of *Gnptab*-mutant mice when compared to control littermates (39 ± 5 % area vs 5 ± 5 % area in control; n = 3 animals per group, *P* = 0.001, unpaired *t*-test; Figure 2).

**Figure 2.**
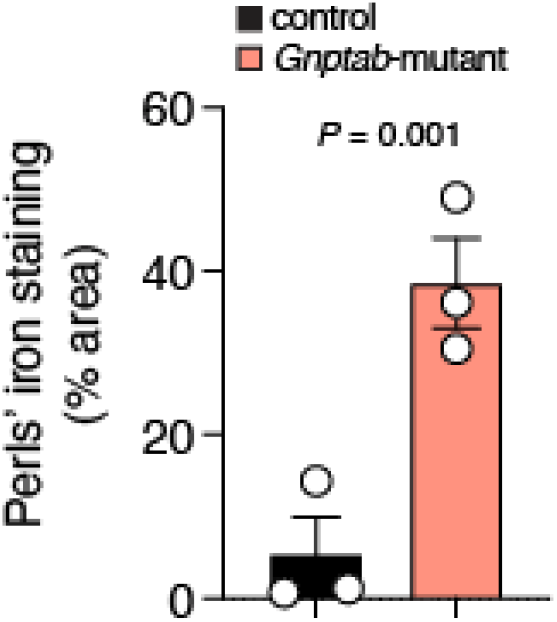
*Gnptab*-mutant mice showed higher iron concentration in the striatum. Summary data shows the percentage area with iron deposition (detected by Perls’ staining) in the striatum reveals a significant increase (∼8-fold) in *Gnptab*-mutant mice compared to controls (unpaired *t*-test, n = 3 animals per group). Data are presented as mean ± sem. *P* – *t*-test

### 3.3 Morphometric characteristics of striatal astrocytes in Gnptab-mutant mice

We and others hypothesized that astrocytes might play a key role in the pathophysiology of stuttering (Han et al., 2019; Maguire et al., 2021; Turk et al., 2021; SheikhBahaei et al., 2023). Therefore, we next evaluated the morphometric properties of striatal astrocytes (Figure 3). Our data suggest that striatal astrocytes are smaller (24073 ± 2573 µm^3^ vs 47507 ± 7762 µm^3^ in control; n = 10 astrocytes, *P* = 0.013, unpaired *t*-test; Figure 3) and less complex (8968 ± 2173 a.u. vs 28134 ± 8099 a.u. in control; n = 10 astrocytes, *P* = 0.032, unpaired *t*-test; Figure 3) when compared to those in the control mice.

**Figure 3.**
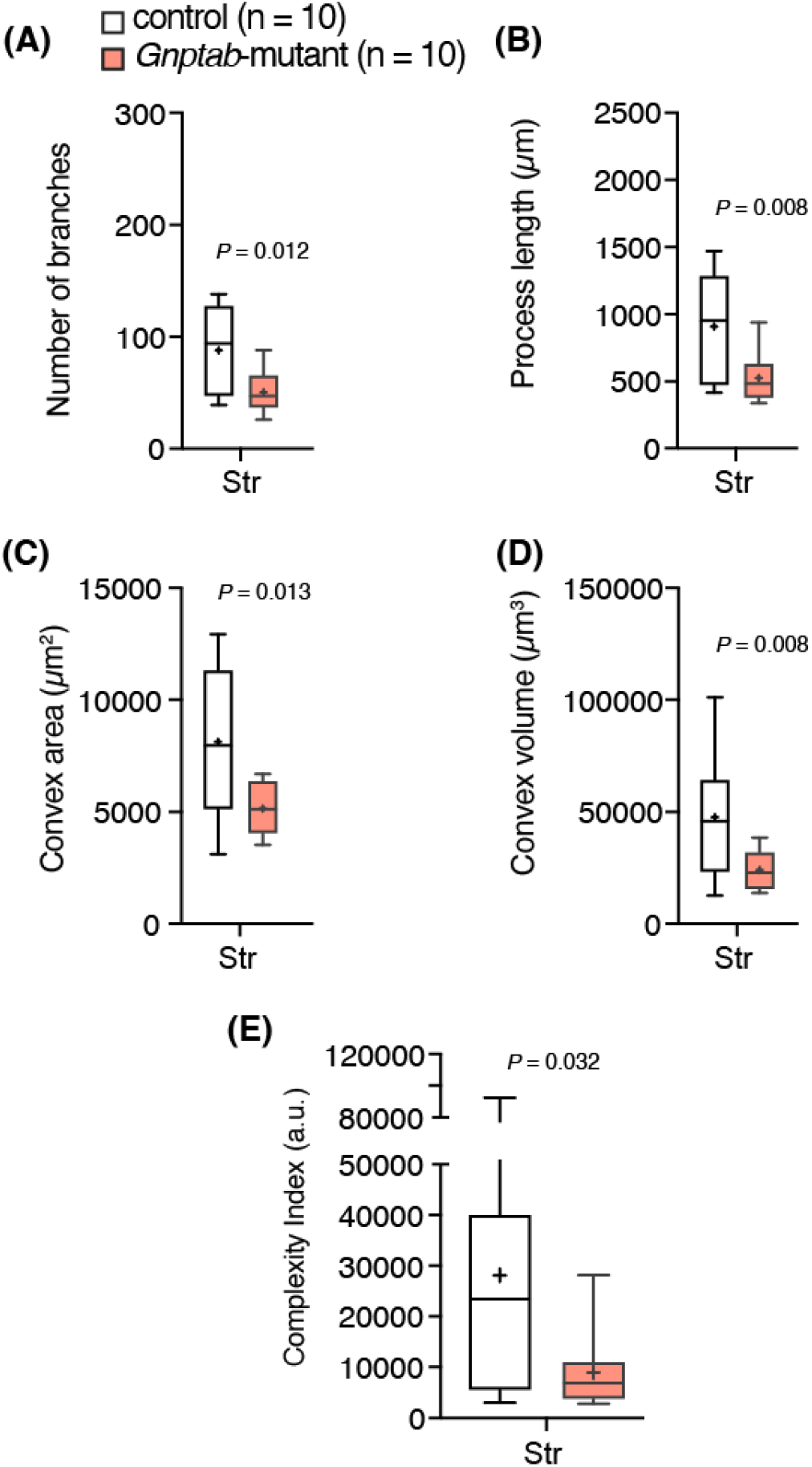
Morphometric analysis of striatal astrocytes in *Gnptab*-mutant and control mice. Summary data illustrating morphometric analysis of striatal astrocytes: (**A**) number of branches, process length, (**C**) convex hull surface area, (**D**) convex hull volume. (**E**) Comparison of normalized data using complexity index (see Methods). Overall, striatal astrocytes were smaller and less complex in *Gnptap*-mutant mice than in control littermates. Str, striatum. n = 10 astrocytes per region from 3 animals per group. *P* – *t*-test

### 3.4 Iron chelator treatment improved vocal behaviors in Gnptab-mutant mice

We next hypothesized that decreasing brain iron levels might improve the behavioral phenotypes in *Gnptab*-mutant mice. Accordingly, we administered an oral iron chelator, DFP, to decrease the iron content in the striatum and analyzed vocalization behaviors (see Methods). DFP (0.1 mg/L in drinking water) improved vocalization phenotypes (i.e., decreased frequency of compound vocalizations) in *Gnptab*-mutant mice (Figure 4).

**Figure 4.**
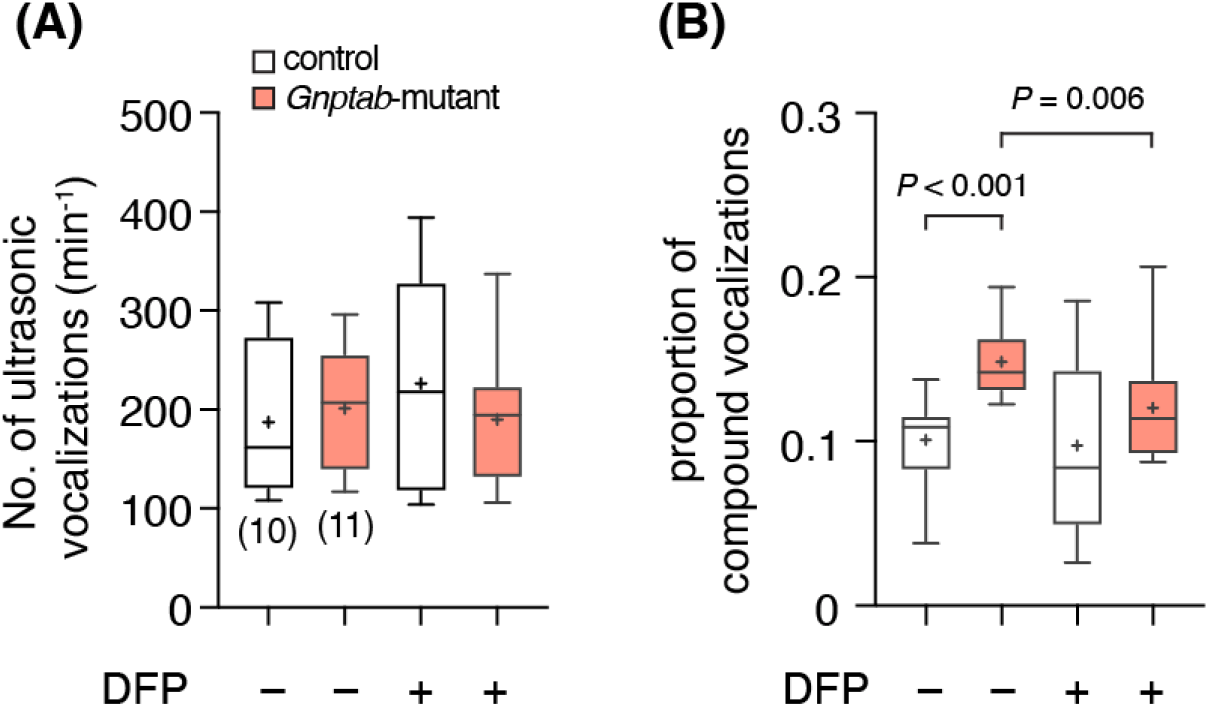
Iron chelation decreased compound vocalizations in *Gnptab*-mutant mice. Summary data showing the effect of iron chelator deferiprone (DFP, 0.1 mg/L in drinking water) on vocalization frequency (**A**) and ratio of compound vocalization (**B**) in *Gnptab*-mutant (n = 11) and control littermates (n = 10). Partially decreased frequency of compound vocalizations indicating improved vocalization phenotypes (*P* – *t*-test).

## 4 DISCUSSION

Childhood-onset fluency disorder, or developmental stuttering, is a neurodevelopmental motor disorder characterized by persistent disruptions in speech fluency (American Psychiatric Association, 2013). Despite being a common speech disorder globally, its etiology remains poorly understood, and no FDA-approved treatments exist to improve the quality of life in people who stutter. In this study, by using a transgenic mouse model with a specific mutation in the *Gnptab* gene linked to stuttering in humans, we showed that iron homeostasis was disrupted in a brain region integral to controlling motor behaviors. Furthermore, we treated the *Gnptab*-mutant mice with the iron chelator drug DFP, which is known to cross the blood-brain barrier (BBB) (Levy et al., 2011). This study showed that oral administration of DFP decreased iron concentration in the brain and improved symptoms of vocalization deficits in *Gnptab-mutant* animals.

### 4.1 Iron accumulation in Gnptab-mutant mice

Previous imaging data from adults who stutter suggested excess iron in the brain regions important for human speech (Cler et al., 2021; Liman et al., 2021). Here, we used histology to *directly* show iron accumulations in the *Gnptab*-mutant mice. Iron-specific Perls’ staining shows a marked increase in iron deposition in *Gnptab*-mutant animals compared to control littermates in adult mice. Our data provide direct evidence for iron deposition in the *Gnptab*-mutant mouse model of stuttering, strengthening correlational findings in humans who stutter (Cler et al., 2021; Liman et al., 2021). The remaining question centers on the definition of a time course of iron accumulation during brain development, if there is one at all, to differentiate this accumulation from that seen in the process of aging and neurodegenerative disorders (Foley et al., 2022; Madden et al., 2023).

### 4.2 Iron homeostasis, lysosomal trafficking dysregulation, and astrocytes

Iron is a necessary transition metal implicated in brain mitochondrial function, myelin production, and the synthesis of neurotransmitters such as dopamine (Ward et al., 2014; Cheli et al., 2020). In the brain, the BBB is responsible for preventing brain access to impermeable compounds such as iron (McCarthy and Kosman et al., 2015; Crichton et al., 2011). Once in the brain, there are complex molecular systems responsible for shuttling iron into cells, storing it, making it biologically available, and, if need be, exporting it from the cell. The coordination of each of these processes, and those yet to be determined, dictates whether iron takes a physiologic or a pathophysiologic role in the central nervous system (Singh et al 2014).

It is evident that properly functioning lysosomes, and lysosomal trafficking pathways, play an important role in cellular iron regulation, though more detail is needed on these processes (Yambire et al., 2019; Kidane et al., 2006; Krishan et al., 2015). For instance, iron that is not immediately used in biological processes is stored in ferritin nanocages in a non-reactive, safe form. This process is dynamic, and when functioning properly, an increased cellular demand for iron will result in lysosomal breakdown of ferritin, delivering biologically active iron to the mitochondria and other cellular iron sinks (Galy et al., 2024). Importantly, acid hydrolases are required for this breakdown of ferritin (Kidane et al., 2006). Therefore, the *Gnptab* gene may play a role in iron dysfunction. The *Gnptab* gene codes for a component of the multi-subunit enzyme that tags acid hydrolases with mannose-6-phosphate (M6P), subsequently directing acid hydrolases to lysosomes. Once acid hydrolases receive a mannose-6-phosphate (M6P) tag they may be recognized by their receptors, initiating clathrin-coated vesicle transport to the lysosome (Le Borgne and Hoflack, 1997; Lin et al., 2004; Alberts et al., 2002; Kang et al., 2019). This trafficking step is essential for establishing lysosomal conditions that allow for ferritin breakdown, suggesting that its partial failure in *Gnptab*-mutant animals may play a role in iron dysregulation.

On the other hand, astrocytes have been implicated to play a critical role in the pathophysiology of developmental stuttering (Han et al., 2019; Turk et al., 2021; Adeck et al., 2024). Astrocytes, proximal to endothelial cells that comprise the BBB, are essential for the process of iron transfer into and around the brain (Dringen et al., 2007). With a deficit in the normal function of lysosomes in *Gnptab*-mutant animals, it is plausible that iron is mainly accumulated in the astrocytes, which may affect their function.

### 4.3 Iron chelation improves vocal phenotype in a mouse model of developmental stuttering

Mouse ultrasonic vocalizations, typically between 30Hz and 120Hz, represent innate (i.e., not learned) behavior most common during pup development (e.g., pup calls) and later related to mating responses (e.g., courtship vocalization in males). Transgenic *Gnptab-mutant* mice have ultrasonic vocal patterns with a decrease in vocalization frequency and an increase in pauses between bouts (Barnes et al., 2016). Our data confirmed that though the frequency of vocalization was similar, the geometry of vocalization was different in adult animals, where the *Gnptab-*mutant mice exhibited more compound vocalizations compared to control animals (Figure 4). Strikingly, administration of the oral iron chelator drug (Deferiprone; DFP), via water supply over one-month, lowered CNS iron levels in brain tissue and improved vocal phenotype in *Gnptab*-mutant male mice. Our data provides a mechanistic explanation in which a mutation in the *GNPTAB* gene, a housekeeping gene involved in cellular trafficking, contributes to an irregularity of vocal behaviors. Equally important, our data further strengthens the evidence supporting the critical role of astrocytes in the pathophysiology of developmental stuttering disorder.

## Acknowledgments

We are grateful for the invaluable mentorship, support, and discussion with Drs. D. Leopold (NIMH), Y. Chudasama (NIMH), and D. Drayna (NIDCD). We thank NINDS Electron Microscopy Core (Director: Dr. J-H Tao-Cheng), Light Microscopy Core (Director: Dr. C. Smith), as well as Drs. S. Bradley (NIMH) and K.F. Messanvi (NIMH) for technical support. We also thank Drs. Dl Reich (NINDS) and S. K. Ha (University of Pittsburgh) for help with Perls’ staining protocol. We especially thank Drs. A.P. Koretsky (NINDS) and T. Rouault (NICHD) for helpful discussions on the manuscript. We also thank Dr. D. Drayna (NIDCD) for providing *Gnptab*-mutant animals. This work was supported by the Intramural Research Program of the NIH, NINDS (NS009420 to SSB), and in part by the NIMH IRP Rodent Behavioral Core (MH002952).

